# Exosomes from adipose-derived stem cells alleviate myocardial infarction via microRNA-31/FIH1/HIF-1α pathway

**DOI:** 10.1101/2021.05.16.444060

**Authors:** Dihan Zhu, Yang Wang, Miracle Thomas, KeAsiah McLaughlin, Babayewa Oguljahan, Joshua Henderson, Qinglin Yang, Y. Eugene Chen, Dong Liu

**Affiliations:** Cardiovascular Research Institute, Morehouse School of Medicine, Atlanta, GA; Department of Pharmacology, Louisiana State University School of Medicine, New Orleans, LA; Department of Internal Medicine, University of Michigan Medical Center, Ann Arbor, MI; Department of Physiology, Morehouse School of Medicine, Atlanta, GA

**Author notes:** **Correspondence:** Dong Liu, M.D., Ph.D., Morehouse School of Medicine, 720 Westview Drive SW, Atlanta, GA 30310. Dihan Zhu and Yang Wang contributed equally to this work.

**Keywords:** stem cell, exosome, angiogenesis, myocardial infarction, microRNA

## Abstract

Our previous study has revealed that exosomes from adipose-derived stem cells (ASCs) promote angiogenesis in subcutaneously transplanted gels by delivery of microRNA-31 (miR-31) which targets factor inhibiting hypoxia-inducible factor-1 (FIH1) in recipient cells. Here we hypothesized that ASC exosomes alleviate ischemic diseases through miR-31/FIH1/hypoxia-inducible factor-1α (HIF-1α) signaling pathway. Exosomes from ASCs were isolated by sequential centrifugations, and were characterized with nanoparticle tracking analysis, transmission electron microscopy, and immunoblotting analysis for exosomal markers. Results from laser imaging of ischemic mouse hindlimb revealed that exosomes enhanced the blood perfusion, and this enhancement was impaired when using miR-31-depleted exosomes. Immunohistochemistry analysis showed that administration of exosomes resulted in a higher arteriole density and larger CD31^+^ area in ischemic hindlimb than miR-31-delpleted exosomes. Similarly, knockdown of miR-31 in exosomes reduced the effects of the exosomes on increasing ventricular fraction shortening and CD31^+^ area, and on decreasing infarct size. Exosomes promoted endothelial cell migration and tube formation. These changes were attenuated when miR-31 was depleted in the exosomes or when FIH1 was overexpressed in the endothelial cells. Furthermore, the results from co-immunoprecipitation and luciferase reporter assay demonstrated that the effects of exosomes on elevating the binding of HIF-1α with co-activator p300 and enhancing HIF-1α activity were decreased when miR-31 was depleted in the exosomes or FIH1 was overexpressed. Our findings provide evidence that exosomes from ASCs promote angiogenesis in both mouse ischemic hindlimb and heart through transport of miR-31 which targets FIH1 and therefore triggers HIF-1α transcriptional activation.

## INTRODUCTION

Ischemic diseases continue to represent a critical and growing cause of morbidity and mortality in modern society[1]. Peripheral artery disease (PAD) is a narrowing or blockage of the blood vessels causing ischemia in limbs[2]. Ischemic heart disease, also known as coronary heart disease, is caused by the stenosis and occlusion of the coronary artery, which leads to insufficient blood supply to the heart [2]. To address these challenges, the science of therapeutic angiogenesis has been developing for the past two decades. The protein/gene approach, stem/progenitor cell approach, and subsequent microvesicle/exosome approach have been extensively studied in the treatment of ischemic diseases[3, 4]. Exosomes are plasma membrane-derived vesicles released by cells and are recognized as transmitters loaded with RNAs, proteins, and lipids that participate in intercellular communication[5]. In comparison with the stem cell approach, the cell-free exosome approach avoids the possibility of tumor formation due to uncontrolled cell proliferation and microvasculature occlusion upon intra-arterial administration of implanted cells[3]. Unlike other stem cells, ASCs possess the characteristics of easy acquisition and less immunogenicity, making them attractive candidates as exosome donors[6]. We are the first to report that exosomes derived from ASCs promote angiogenesis in animals[7] which is subsequently confirmed by others[8–10]. However, the underlying molecular mechanisms remain largely unclear, thus hindering their clinical translation.

HIF-1 plays a crucial role in adaptive responses to hypoxia/ischemia by driving transactivation of dozens of genes involved in angiogenesis[11, 12]. HIF-1 is a heterodimeric transcription factor composed of oxygen-regulated HIF-1α and constitutively expressed HIF-1βsubunits. Hydroxylation of asparagine residues of HIF-1α by FIH1 as a post-translational modification prevents HIF-1α binding to the transcriptional co-activator p300 and therefore abrogates the subsequent HIF-1α-mediated gene transcription[13]. It has been well established that inhibition of FIH1 releases HIF-1α from inactivation and promotes angiogenesis[14, 15]. The microRNAs (miRNAs) are small, non-coding RNAs that regulate various biological processes, including angiogenesis[16, 17]. Our studies, as well as the research of others, has shown that miR-31 targets FIH1 and thus inhibits its expression[7, 18, 19]. The majority of RNA in stem cell-derived exosomes is miRNA, which contributes to exosome-based therapy for ischemic diseases[7, 20–22]. In this study, we demonstrate that exosomes from ASCs enhance angiogenesis in mouse ischemic hindlimb and heart via the delivery of miR-31, which targets FIH1 and consequently initiates HIF-1α transactivation.

## MATERIALS AND METHODS

### Cell culture

Human adipose-derived stem cells (ASCs) and human microvascular endothelial cells (HMVEC) were purchased from Thermo Fisher Scientific (Carlsbad, CA). Both cell types were maintained at 37°C in a humidified 5% CO_2_ incubator. Cells of passages 4-6 were used for all experiments.

### Exosome isolation

ASCs were preconditioned with endothelial differentiation medium (PromoCell, Heidelberg, Germany) for 4 days as described previously[7]. Exosomes were isolated by sequential centrifugations of the ASC culture medium at 500 × g for 10 minutes, 12,000 × g for 30 minutes, and 100,000 × g for 60 minutes at 4°C. To remove residual soluble factors, pelleted exosomes were then washed with PBS once by centrifugation at 100,000 × g for 60 minutes at 4°C. The protein concentration of the isolated exosomes was detected by using a total exosome RNA and Protein Isolation Kit (Thermo Fisher Scientific) and a BCA Protein Assay Kit (Thermo Fisher Scientific) according to the manufacturer’s instructions. In all cases, the FBS used in this study was exosome-free FBS, which was obtained by ultracentrifugation of diluted FBS at 100,000 × g for 18 hours at 4°C.

### Nanoparticle tracking analysis

A Nanosight LM10 instrument (Malvern; Amesbury, UK) was used to perform nanoparticle tracking analysis according to the manufacturer’s recommendations. A total of 500 μl of a 1:5-diluted exosome sample in phosphate-buffered saline (PBS) was injected into a NanoSight sample cubicle. Each exosome sample was analyzed by detecting the rate of the Brownian motion of particles in liquid suspension. The analysis settings were optimized, and each video was analyzed to obtain the mean, mode, median, and estimated concentration of each particle size.

### Transmission electron microscopy

Transmission electron microscopy of isolated exosomes was performed at the Integrated Electron Microscopy Core, University of Emory. Briefly, freshly-isolated exosome samples were fixed in 2.5% glutaraldehyde and dried on 300 mesh copper grids. Exosomal specimens were then negatively stained with 2% aqueous uranyl acetate and visualized using a JEOL JEM-1400 transmission electron microscope (Tokyo, Japan) equipped with a Gatan BioScan US1000 CCD camera (Gatan, Inc., Pleasanton, CA).

### Immunoblotting

Immunoblotting was performed as previously described[7]. Proteins were detected using primary anti-CD9 (ab92726, Abcam), anti-TSG101 (T5701, Millipore Sigma), anti-FIH1 (SAB2101040, Millipore Sigma) and anti-GAPDH (437000, Thermo Fisher Scientific) antibodies. Exposure of the resultant protein bands was performed with an ImageQuant LAS 4000 Luminescent Image Analyzer (GE Healthcare; Chicago, IL).

### Creation and transduction of recombinant lentivirus

The procedures performed here followed the National Institutes of Health guidelines for recombinant DNA research. To knockdown the expression of miR-31 in ASCs, a lentiviral Lenti/anti-miR-31 (antimiR-31) was created using commercial plasmids pLenti/anti-miR-31 (System Biosciences, Palo Alto, CA). A Lenti/anti-miR-Cont (antimiR-Cont) was created as a corresponding control. To overexpression of FIH1 in HMVECs, a lentiviral Lenti/FIH1 was purchased from Amsbio (Abingdon, UK). A lentiviral Lenti/Cont was purchased as a corresponding control. The detailed methods of lentiviral creation, concentration, titration, and transduction were described previously[7, 23].

### RNA isolation and RT-PCR for miRNAs

Total RNA was extracted from exosomes using TRIzol Reagent (Thermo Fisher Scientific) according to the manufacturer’s instructions. Reverse transcription and quantitative real-time PCR were performed using the TaqMan miRNA assay system (Thermo Fisher Scientific) according to the manufacturer’s instructions and our previous description[7]. The relative miRNA levels were normalized to endogenous U6 small nuclear RNA for each sample.

### Hindlimb ischemia (HLI) model

All animal experiments in this study were approved by the Institutional Animal Care and Use Committee of the Atlanta University Center and complied with the NIH guidelines for the care and use of laboratory animals. The femoral artery was ligated on the left hindlimb of male C57BL/6J mice (The Jackson Laboratory, Bar Harbor, ME) at age 6-8 weeks according to our previous report[24]. The mice were subjected to intramuscular injection in the left adductor muscle with PBS or 30 μg of various exosomes from ASCs immediately after ligation. The same intramuscular injections were performed at the surgery site 4 and 8 days post-surgery. Blood perfusion on the plantar surface of the hind paws was measured immediately before and after surgery and 3, 7, 14, and 21 days post-surgery using laser speckle imaging (LSI, Perimed AB, Järfälla, Sweden). The left to right ratio (L/R) was used to represent the relative blood perfusion rate at the left hindlimb for each animal (n = 10). The mice were euthanized 21 days after the surgery. The left adductor muscle and gastrocnemius muscle were collected and cryosectioned for immunohistochemistry analysis as described below.

### Myocardial infarction (MI) model

C57BL/6J mice (10-12 weeks old) were kept anesthetized on a heating pad in a supine position by inhalation of 2% isoflurane via a face mask. Surgery procedures were performed as Gao et al described[25]. Briefly, a skin cut, 10-12 mm in length, was made over the left chest, and a purse-string suture was made around the incision. The fourth intercostal space was opened after dissection and retraction of the pectoral major and minor muscle. The heart was immediately and gently “popped out” through the opening. The left anterior descending (LAD) coronary artery was ligated. PBS or various exosomes from ASCs was intramyocardially injected in the infarct border area (5 μg per injection for two injections on each side of the ligation) by using a 32 G microsyringe (Hamilton, Reno, NV). The heart was allowed to retract in the thoracic cavity followed by manual evacuation of air and closure of the skin opening with the previously placed purse-string suture. Intravenous injection of PBS or 100 μg of various exosomes from ASCs was performed through tail vein at 7, 14, 21 days post-surgery. The mice were euthanized 28 days post-surgery. Hearts were collected, paraffin-embedded, transversely sectioned, and subject to Masson’s staining and immunohistochemistry analyses as described below.

### Echocardiography

Echocardiography was performed twice for each mouse at 1 day before and 28 days after the LAD coronary artery ligation. Mice were anesthetized with inhalation of isoflurane. Transthoracic 2-dimensional images were obtained using a high-resolution echocardiography system HP5500 (Hewlett-Packard, San Jose, CA) equipped with a 15-MHz transducer. Two-dimensional guided M-mode was then used to measure left ventricle (LV) end-systolic diameter (LVESD) and LV end-diastolic diameter (LVEDD) at the mid-ventricular level. The percentage of LV fractional shortening (LVFS%) was calculated as ((LVEDD−LVESD)/LVEDD×100%).

### Masson’s staining

MI was evaluated by using a trichrome stain (Masson) kit (Millipore Sigma Aldrich) on cardiac sections according to the manufacturer’s instructions. The normal myocardium showed red while the infarcted myocardium showed blue due to a collagen fibril formation. Infarct sizes of LV were evaluated as described in a previous report[26]. Briefly, a LV myocardial midline was drawn at the middle between the epicardial and endocardial surfaces by using an ImageJ software (Media Cybernetics). The percentage of the arc length occupied by infarcted myocardium in the LV circumference was calculated as the infarct sizes of the myocardial section. The average percentage from four layers at 1 mm intervals was used to represent the infarct sizes of LV.

### Immunohistochemistry

The muscle sections were stained with the primary anti-CD31 antibody (1:500, 550274, BD Biosciences) and counterstained with DAPI as described previously[24]. The regions containing the most intense CD31^+^ areas of neovascularization were chosen for quantification. Five hotspots per section and 3 sections per muscle were analyzed at 400x magnification. ImageJ software was used to measure CD31^+^ areas in each hotspot. The adductor muscle was additionally incubated with the primary anti-α-smooth muscle actin (α-SMA, 1:500, 19245, Cell Signaling, Danvers, MA). Arteriole density was assessed as α-SMA^+^ vessels/mm^2^.

### Exosome uptake and flow cytometry analysis

Exosome uptake and flow cytometry analysis was performed as described previously[24]. In brief, exosomes from ASCs were labeled fluorescent red using an ExoGlow-Protein EV Labeling Kit (System Biosciences) and then incubated with precipitation solution at 4°C overnight. The labeled exosomes were precipitated by centrifugation at 10,000 rpm for 10 minutes and resuspended in HMVEC basal medium. HMVECs were marked green by incubating in 5 μmol/L cell-permeable fluorescein, carboxyfluorescein succinimidyl ester (Thermo Fisher Scientific) at 37°C for 20 minutes. The cells were washed and then incubated with the labeled exosome suspension at a concentration of 50 μg/ml for the indicated times (0, 6, 12, or 24 hours) or at the indicated concentrations (0, 25, 50, or 100 μg/ml) for 24 hours. The cells were washed and then incubated with fresh medium containing 100 ng/ml Hoechst 33342 (Thermo Fisher Scientific) at 37°C for 30 minutes. Olympus DP-70 fluorescent microscopy (Olympus, Japan) was used to visualize the uptake of exosomes to HMVECs, and the images were analyzed by using an Image-pro plus 6.2 software. For flow cytometry analysis, the cells were collected and analyzed by using a Guava easyCyte flow cytometer equipped with CytoSoft software (Millipore Sigma). ExpressPlus program was selected. The gain controls were kept at their default settings. Each cell was measured for forward scatter (FSC-HLog) versus Red fluorescence (RED-HLog). The results were further analyzed by using FlowJo software.

### Migration assay

HMVECs were treated with various exosomes at 50 μg/ml (protein concentration) for 24 hours with or without a pre-transduction of Lenti/Cont or Lenti/FIH1 as described above. The treated HMVECs were then seeded in a 24-well insert plate (BD Biosciences; San Jose, CA) and incubated at 37°C for 24 hours. Cell nuclei were stained with Hoechst 33342 (Thermo Fisher Scientific). The cells that migrated to the lower side of the inserts were counted as described in our previous report[24].

### Tube formation assay

HMVECs were treated with various exosomes at 50 μg/ml (protein concentration) for 24 hours with or without a pre-transduction of Lenti/Cont or Lenti/FIH1 as described above. The treated HMVECs were seeded on precoated growth factor-reduced Matrigel (BD Biosciences) and incubated with endothelial basal medium/1% FBS at 37°C for 4 hours. The cells were then stained with Calcein AM (Thermo Fisher Scientific), and tube formation was visualized using fluorescence microscopy. The total vessel length, vessel area, number of junctions, and network complexity were calculated using AngioTool v.2 software[24].

### Co-immunoprecipitation

Protein A magnetic beads (Cell Signaling Technology) were pre-washed with cell lysis buffer twice. Magnetic separation rack (Cell Signaling Technology) was used to separate magnetic beads from liquid buffer. HMVEC nuclei were isolated by using NE-PER Nuclear and Cytoplasmic Extraction Reagents (Thermo Fisher Scientific). The nuclear lysate was pre-cleared by incubating with the pre-washed magnetic beads for 20 min at room temperature with rotation. The pre-cleared nuclear lysate was incubated with anti-p300 primary antibody (70088, Cell Signaling Technology) overnight at 4°C with rotation to form immunocomplex which was subsequently incubated with the pre-washed magnetic beads for 20 min at room temperature with rotation. Normal IgG was used parallelly as an isotype control for anti-p300 antibody. The bead-bound immunocomplex was washed five times with lysis buffer, resuspended in loading buffer and heated at 95°C for 5 min to unbind the beads. The free beads were removed from samples by using magnetic separation rack. The samples were subject to immunoblotting with anti-HIF-1α antibody (36169, Cell Signaling Technology).

### HIF-1 activation assay

The eLUCidate HRE reporter cell line, a stably transfected HeLa cell line which expresses Renilla luciferase reporter gene under the transcriptional control of the hypoxia response element (HRE), was obtained from Genlantis (San Diego, CA). The cells were seeded in a 96-well plate at 1×10^4^ cells per well and treated with various exosomes at 50 μg/ml for 48 hours. The HRE signals were detected by using a Renilla Luciferase Glow Assay Kit (Thermo Fisher Scientific). Briefly, the cells were rinsed with PBS once. Cell lysis buffer was added at 50 μl/well followed by shaking on a shaker for 15 min. For each well, 20 μl of the cell lysate was transferred to an opaque 96-well plate and incubated with 50 μl of working solution for 10 min. The light output was detected with a SpectraMax M5 microplate reader (Molecular Devices, San Jose, CA).

### Statistical analysis

Statistical significance between two groups was evaluated with a two-tailed Student’s *t* test. For multiple group comparisons, one-way ANOVA followed by Tukey’s multiple comparison test were performed. All values were reported as the mean ± SD. A difference was considered significant when *p*<0.05.

## Results

### MiR-31 contributes to the proangiogenic effects of exosomes from ASCs in mouse ischemic extremity and heart

Exosomes are small particles secreted from cells at sizes ranging from 30-150 nm[27]. In this study, nanoparticle tracking analysis was performed to determine the size distribution profile and relative particle density. Particles that collected from ASC medium were exhibited within the range of exosomes (Figure 1A), in agreement with our previous report[24]. In addition, we performed single vesicle analysis for visualization at high resolution with transmission electron microscopy which exhibited typical morphologic characteristics of exosomes, appearing as submicron vesicles lacking a nucleus with a condensed core encapsulated by membrane (Figure 1B). Both transmembrane protein marker (CD9) and cytosolic protein marker (TSG-101) of exosomes were detected in our isolated particles. Furthermore, the results demonstrated that the preconditioning of exosomal donor cells with endothelial cell differentiation medium augmented the harvest of exosomes (Figure 1C). All together, these data show that our isolated exosomes from ASCs are in accordance with the minimal information for studies of extracellular vesicles 2018 (MISEV2018) guidelines[28].

**Figure 1.**
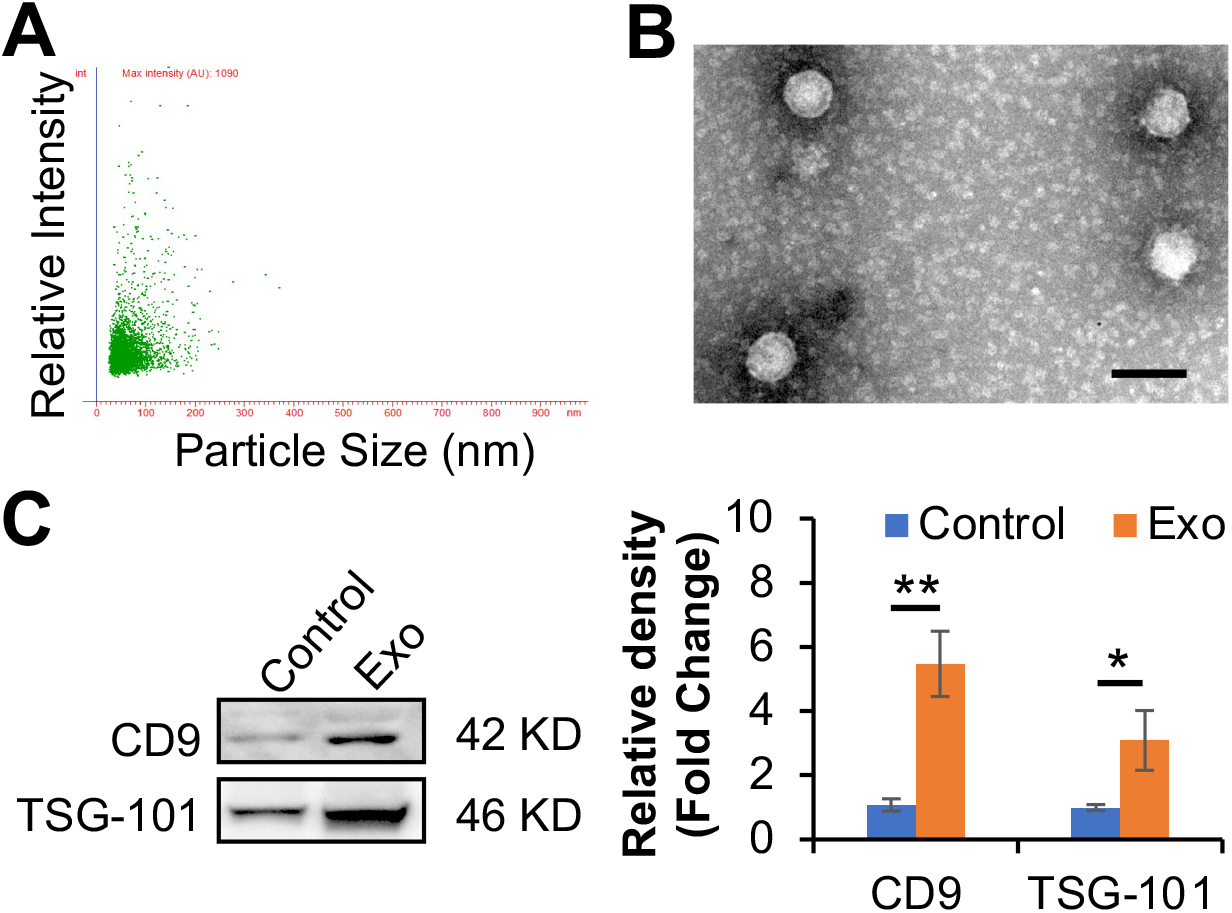
Characterization of exosomes from ASCs. **A**, The isolated exosomes were examined using nanoparticle tracking analysis. Scatter plot graphs of exosomes demonstrated the particle size versus light intensity of exosomes. **B**, Morphology of exosomes was visualized under transmission electron microscopy (scar bar: 100nm). **C**, Exosomal markers, CD9 and TSG101, in exosomes from ASCs with endothelial differentiation medium preconditioning (Exo) were determined via immunoblotting analysis. Exosomes from ASCs without preconditioning were used as a control. Each lane represented an exosomal lysate collected from 1 × 10^6^ ASCs. The right panel is a statistical analysis of the density of the immunoblotting bands using Image J software (n = 4). Protein levels of the control were set to 1. \**p*<0.05, \*\**p*<0.01.

Our previous studies have demonstrated that miR-31 is enriched in exosomes from ASCs and contributes to exosomal proangiogenesis in subcutaneously transplanted gel plugs[7]. To explore the therapeutic angiogenesis in PAD, a common disorder and a major cause of morbidity and mortality worldwide[29], we here evaluated the angiogenic effect of the exosomal miR-31 in a mouse HLI model. Firstly, miR-31-downregulated exosomes were acquired from ASCs that transduced with a lentiviral antimiR-31 (Figure 2A). Then the exosomes with or without miR-31-downregulation were injected intramuscularly into HLI mice. Exosomes from ASCs that transduced with a lentiviral antimiR-control or PBS were used as a control, respectively. Our results from non-invasive laser speckle images revealed that the blood perfusion in the hind paw in mice that were administered normal exosomes exhibited better recovery than those that were administered PBS starting from day 3 (*p* = 0.024) post femoral artery ligation, while the blood perfusion in mice that were administered miR-31-downregulated exosomes exhibited less recovery than those that were administered the corresponding control starting from day 14 (*p* = 0.004) post femoral artery ligation (Figure 2B and 2C). The exosomes enhanced the recovery of blood perfusion by 40.3% in comparison with PBS at three weeks post-surgery (Figure 2D). The enhancement was significantly impaired when using miR-31 depleted exosomes. The results of immunohistochemistry analysis showed that the vascular endothelial cell marker, CD31, positive area in the adductor muscle (Figure 2E and 2F) and gastrocnemius muscle (Figure 2G and 2H) was elevated by a treatment of exosomes. The elevation was reduced when miR-31 was downregulated in the exosomes. As expected, the administration of exosomes was observed to make an increase in arteriole density in adductor muscle, which was suppressed when miR-31 was downregulated in the exosomes. These data suggest that exosomal miR-31 promotes angiogenesis and arteriogenesis in mouse ischemic hindlimb.

**Figure 2.**
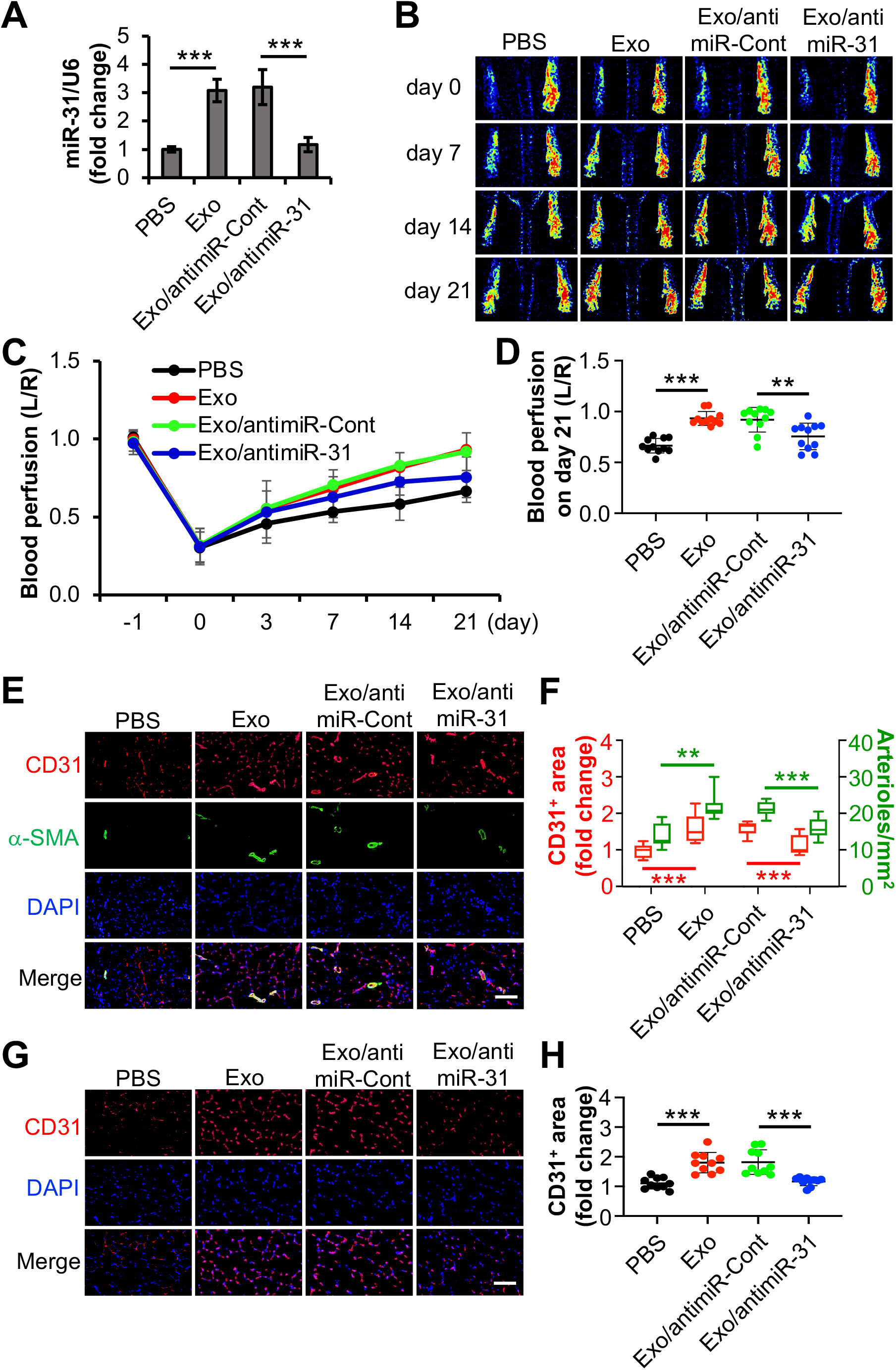
miR-31 contributes to the angiogenic effects of exosomes from ASCs on mouse HLI model. **A**, Validation of lentiviral anti-miR-31 to knockdown miR-31. Exo or exosomes from transduced ASCs with lentiviral anti-miR-31 (Exo/antimiR-31) were used to treat HMVECs (n = 4). PBS and exosomes from ASCs transduced with antimiR-Cont (Exo/antimiR-Cont) were used as controls. The miR-31 content in the HMVECs was determined with RT-PCR. U6 snRNA was used as an internal control. **B-H**, After a ligation of the left femoral artery in mice, PBS, Exo, or Exo/antimiR-31 were intramuscularly injected in adductor muscle, respectively (n = 10). Exo/antimiR-Cont was used as a control. **B**, Representative laser speckle images showed the recovery of blood perfusion in the hind paws on the indicated days. Day 0 represents the perfusion immediately after the femoral artery ligation. **C and D**, Quantitative analyses of the images on panel B showing the left/right ratio of plantar perfusion. **E-H**, The mice were euthanized three weeks post-surgery. The sections of the adductor muscle from the ligated side were subjected to immunohistochemistry analysis of CD31, an endothelial cell marker, and α-SMA, a smooth muscle marker. DAPI was used as a nuclear counterstain. (E, scale bar: 100 μm). Quantification of the CD31^+^ area and the arteriole number per mm^2^ on panel E (F). The sections of the gastrocnemius muscle from the ligated side were subjected to immunohistochemistry analysis of CD31 and counterstained with DAPI (G, scale bar: 100 μm). Quantification of the CD31^+^ area (H). The CD31^+^ area on the slide from the mouse administered PBS was set to 1. \*\**p*<0.01, \*\*\**p*<0.001.

To further explore the angiogenic effect of ASC exosomes and the underlying mechanism on ischemic heart disease, we evaluated their effect on a mouse MI model. Four weeks post ligation of LAD coronary artery, the data of echocardiography demonstrated that the percentage of left ventricular fractional shortening (LVFS) was higher in mice that were administered exosomes in comparison with those that were administered PBS, while the percentage in mice administered miR-31-depleted exosomes was lower than the corresponding control (Figure 3A and sFigure 1). The results from the Masson’s staining of the harvested hearts revealed that the percentage of LV infarct size in mice that received exosome treatment was lower. The beneficial effect was partially reversed when miR-31 was depleted from the exosomes (Figure 3B). Meanwhile, the results of immunohistochemistry analysis showed that the CD31 positive area was elevated in the mouse heart which received exosomes (Figure 3C and 3D). Similarly, the elevation was reduced in the heart of mouse which received miR-31-downregulated exosomes. These findings suggest that ASC exosomes improve cardiac function, reduce infarct size, and promote angiogenesis in mouse infarcted heart via delivery of miR-31.

**Figure 3.**
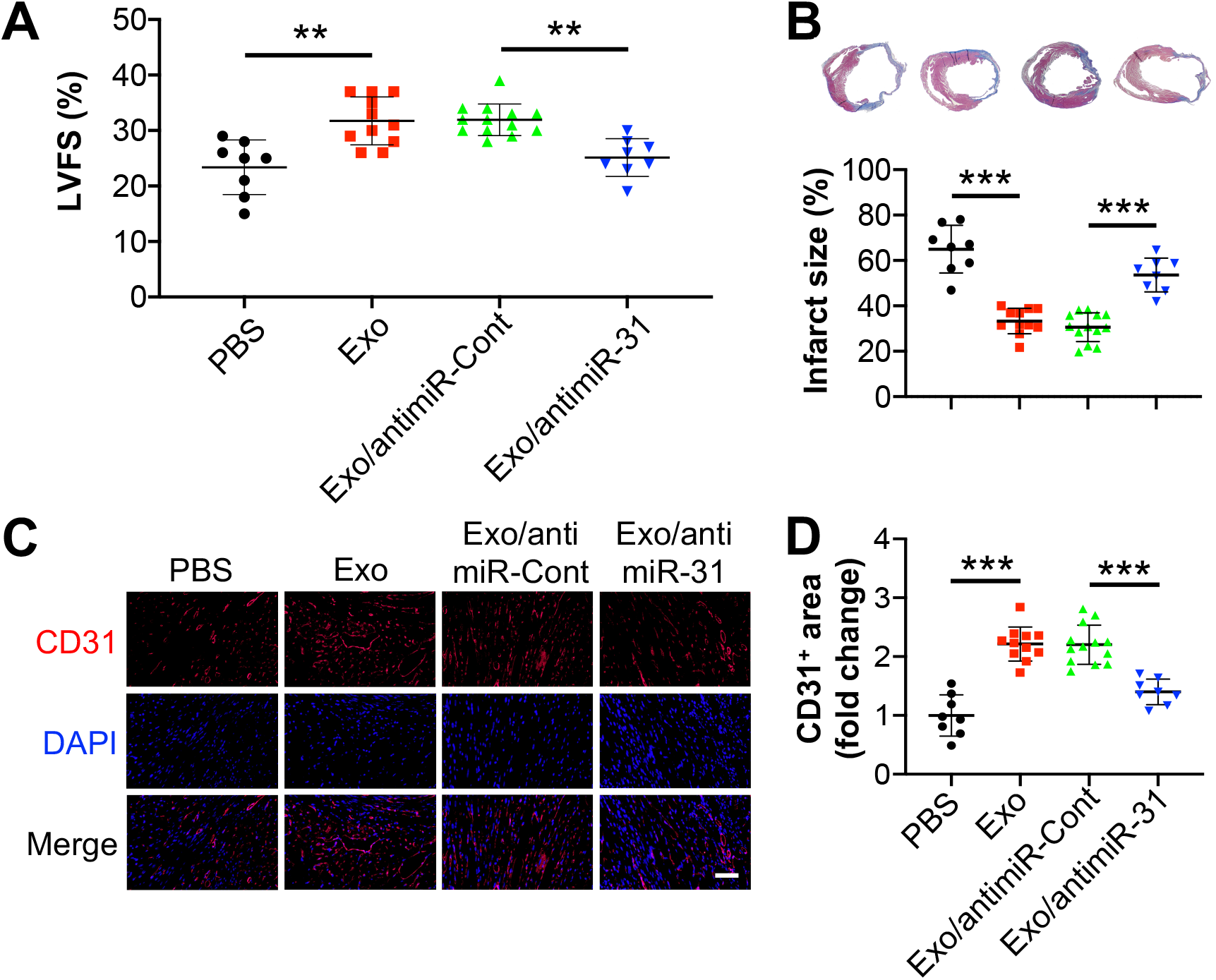
Exosomes from ASCs promote angiogenesis and function recovery in ischemic heart via delivery of miR-31. Mouse myocardial infarction (MI) was induced by permanently ligating the left anterior descending (LAD) coronary artery. PBS (n = 8), Exo (n =11) or exosomes from ASCs transduced with anti-miR-Cont (n = 13) or anti-miR-31 (n =8) were intramyocardial injected, respectively. Four weeks post-surgery, cardiac function was evaluated with echocardiography, and hearts were collected thereafter for morphological examination. **A**, Infarct size was evaluated with fibrotic area of left ventricle by using Masson’s trichrome staining. The blue color represents fibrosis. **B**, Left ventricular fractional shortening (LVFS) was determined with echocardiography. **C**, The sections of the hearts were subjected to immunohistochemistry analysis of CD31, an endothelial cell marker, and counterstained with DAPI (scale bar: 50 μm). **D**, Quantification of the CD31^+^ area. The CD31^+^ area in the PBS group was set to 1. \*\**p*<0.01 and \*\*\**p*<0.001.

### Exosomal miR-31 promotes endothelial cell migration and tube formation by targeting FIH1

Our previous study has both demonstrated that miR-31 contributes to ASC exosomal proangiogenesis in human umbilical vein endothelial cells, and also implied that FIH1 might be a target protein[7]. In this study, we investigated the effect of FIH1 on ASC exosome-induced angiogenesis in HMVECs. Firstly, fluorescence labeled exosomes were used to examine the uptake of exosomes into HMVECs. The results from flow cytometry analysis and fluorescent images showed that the percentage of exosomal internalization increased in a time-course manner and reached 94.4% at 24 hours (Figure 4A and 4B). The percentage of exosomal internalization also increased in a dose-dependent manner and reached 93.6% and 94.5% at the exosomal protein concentrations of 50 μg/ml and 100 μg/ml, respectively (Figure 4C and 4D). An exosomal protein concentration of 50 μg/ml and treatment duration of 24 hours were used for the following *in vitro* experiments. The suppression of FIH1 in HMVECs by treatment of exosomes or transfection of a mimic miR-31 was validated with immunoblotting. Meanwhile, the upregulation of FIH1 levels in HMVECs by transduction of lentiviral FIH1 was also validated. Interestingly, exosome-induced downregulation of FIH1 was attenuated when miR-31 in exosomes was depleted (Figure 5A). The results of HMVEC migration assay showed that exosomes promoted cell migration, which was reversed when miR-31 was depleted in exosomes or when overexpression of FIH1 was accompanied (Figure 5B and 5C). As expected, the exosomes increased HMVEC tube length (Figure 5D and 5E), tube junctions (sFigure 2A), vessel area (sFigure 2B) and decreased network complexing (sFigure 2C). These proangiogenic effects were weakened when miR-31 was depleted in exosomes or when overexpression of FIH1 was accompanied. Our findings provide evidence that miR-31 contributes to exosomal proangiogenesis *in vitro* through targeting FIH1.

**Figure 4.**
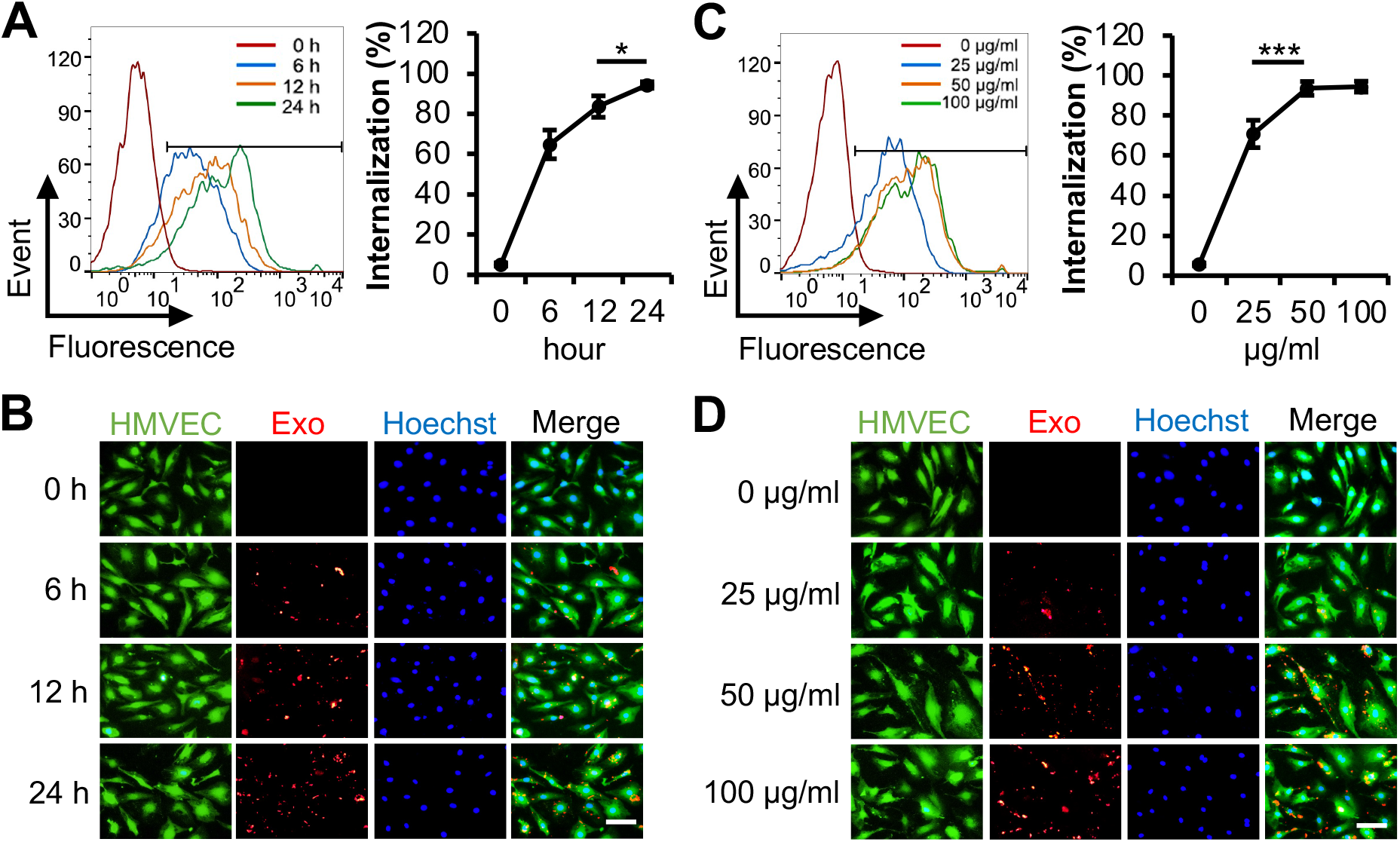
Uptake of the exosomes from ASCs into HMVECs. **A** and **B**, HMVECs were incubated with red-labeled Exo at a concentration of 50 μg/ml for the indicated times. **C and D**, HMVECs were incubated with red-labeled Exo at the indicated concentrations for 24 hours. A and C, The percentage of exosomal internalization into HMVECs were measured using flow cytometry. B and D, Fluorescent images showed the uptake of labeled Exo in HMVECs. Merged images display HMVECs (green) for cell cytoplasm, Hoechst 33342 (blue) for nuclei and red-labeled Exo (n=4). Scale bar: 100 μm. \**p*<0.05, \*\*\**p*<0.001, NS, not significant.

**Figure 5.**
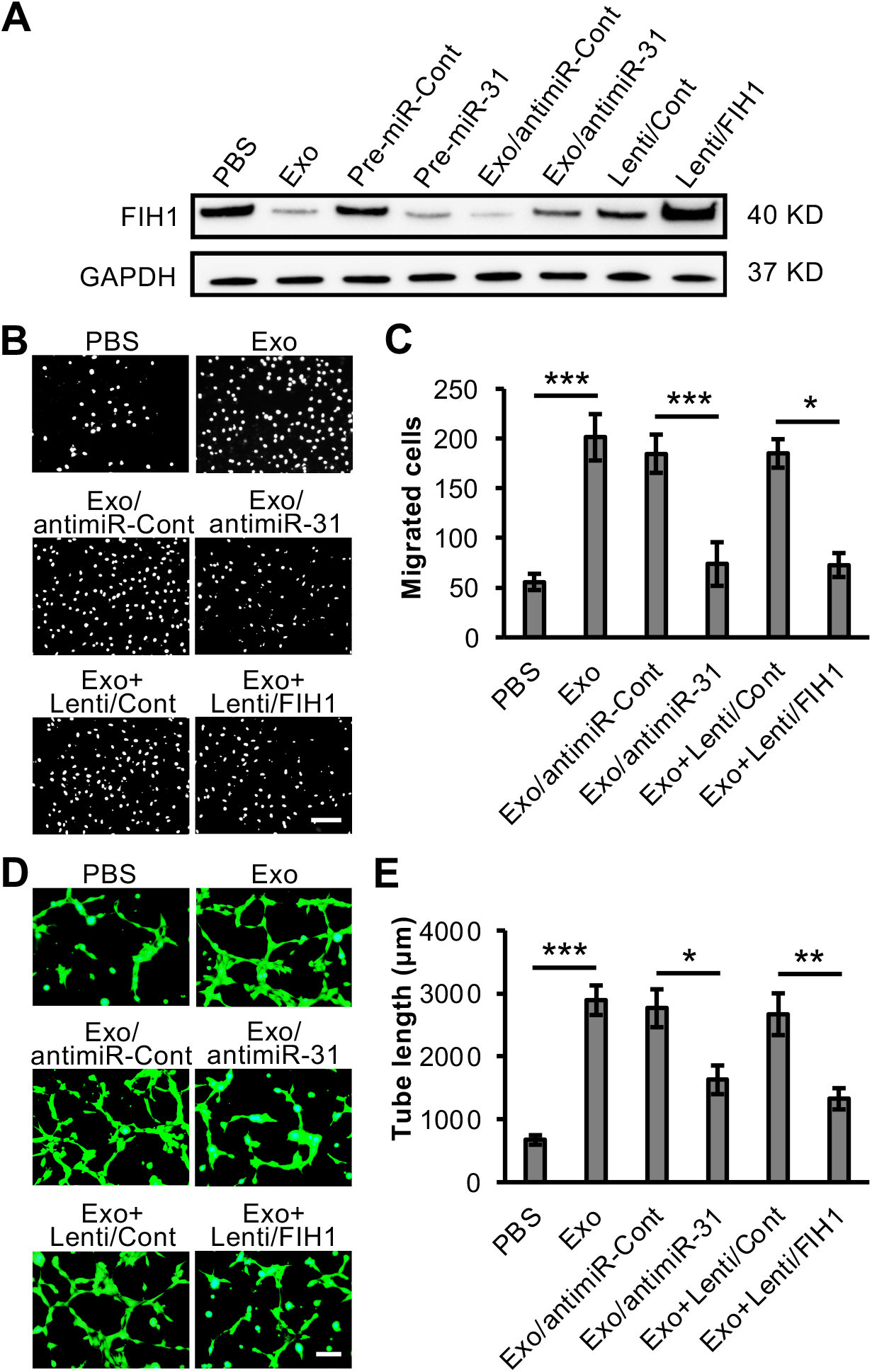
Exosomal miR-31 promotes angiogenesis via suppressing FIH1 in HMVECs. **A**, PBS, Exo or exosomes from transduced ASCs with lentiviral anti-miR-31 (Exo/antimiR-31) were used to treat HMVECs. Exosomes from ASCs transduced with anti-miR-Cont was used as a control (Exo/antimiR-Cont). The levels of FIH1 in HMVECs were determined with immunoblotting. The levels of FIH1 were also examined in HMVECs when overexpression of pre-miR-31 or Lenti/FIH1. Pre-miR-Cont or Lenti/Cont was used as a control, respectively. GAPDH was used as a loading control. **B-E**, HMVECs were treated with Exo, Exo/antimiR-31, or Exo plus lentiviral FIH1 (Exo+Lenti/FIH1). PBS, Exo/antimiR-Cont, or Exo+Lenti/Cont was used as a control, respectively. **B**, Cell migration assay for the treated HMVECs (n = 4). Scale bar: 100 μm. **C**, Quantification of the migrated cells in panel B. **D**, Cell tube formation assay for the treated HMVECs (n = 4). Scale bar: 50 μm. **E**, Quantification of the tube length in panel D. \**p*<0.05, \*\**p*<0.01 and \*\*\**p*<0.001.

### Downregulated FIH1 by exosomal miR-31 increases HIF-1α activity

FIH1 is an asparaginyl hydroxylase that hydroxylates HIF-1α and therefore blocks its binding to p300. Without the binding of co-activator p300, HIF-1α is unable to interact with HRE and to promote angiogenic factor transcription[11, 30]. To investigate the effect of FIH1 and exosomal miR-31 on HIF-1α, we examined the HIF-1α activity with co-immunoprecipitation analysis and luciferase reporter assay. Co-immunoprecipitation system was tested first, which excluded the contamination from beads and anti-p300 antibody and confirmed the specificity of anti-p300 antibody and anti- HIF-1α antibody (sFigure 3). Then the nuclear lysates from HMVECs upon indicated treatments were subject to co-immunoprecipitation. The results showed that although the level of pan HIF-1α in HMVECs remained constant, its level in nuclei, which represents the activation of HIF-1α, was increased upon the treatment of exosomes. This increase was reversed when miR-31 in exosomes was depleted (Figure 6A). Moreover, overexpression of FIH1 also reversed the exosomes-induced increase of p300-bound HIF-1α level (Figure 6B). In addition, similar effects on HIF-1α activity were observed with a stable HeLa cell line expressing a HIF-1α-activated luciferase reporter gene (Figure 6C and 6D). These data suggest that the miR-31-FIH1-HIF-1α signaling pathway may mediate ASC exosome-induced angiogenesis.

**Figure 6.**
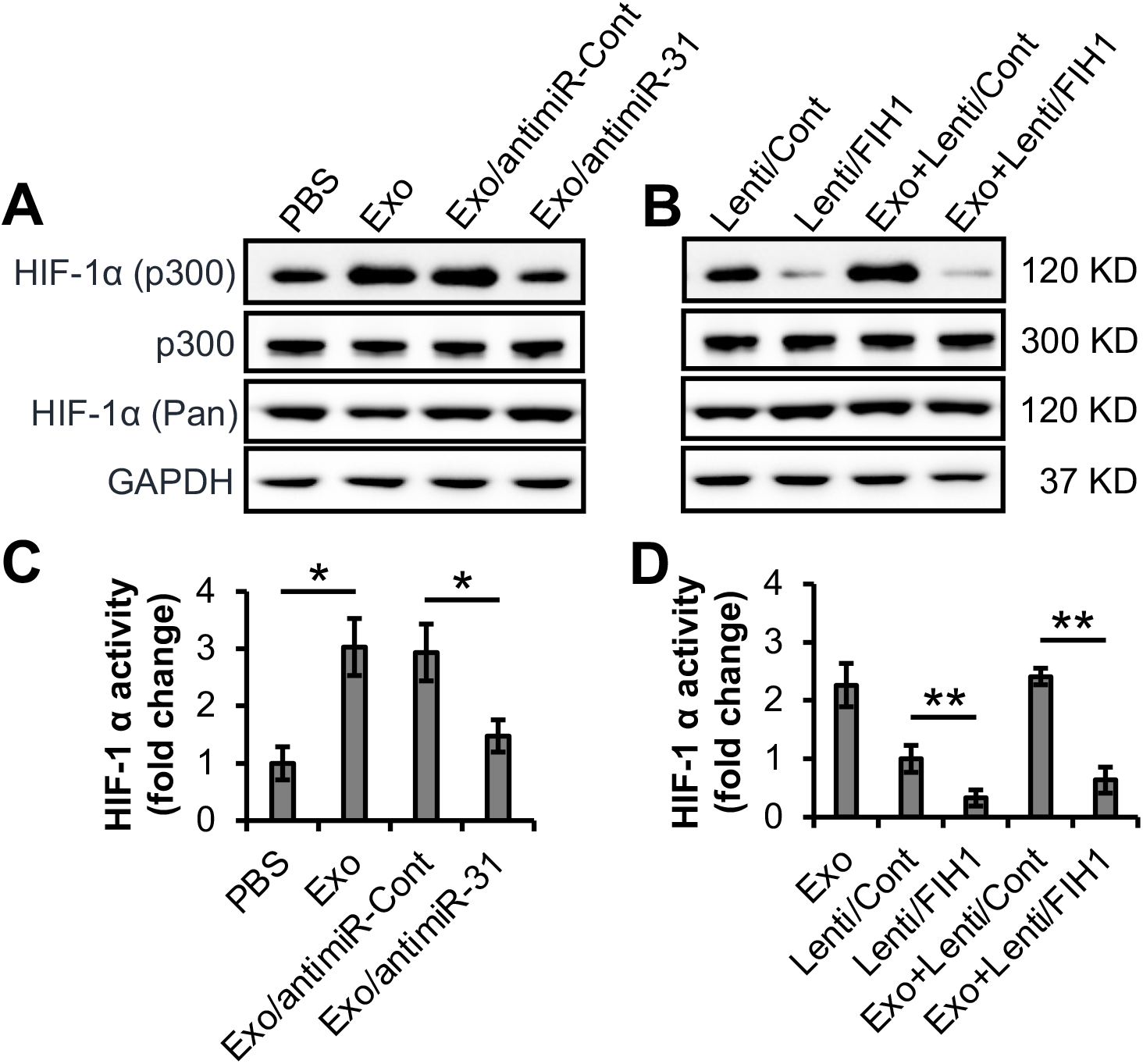
Exosomal miR-31 activates HIF-1α via suppressing FIH1. **A and B**, HMVECs were treated with Exo or Exo/antimiR-31. PBS or Exo/antimiR-Cont were used as a control, respectively (A). The cells were also treated with lentiviral FIH1 (Lenti/FIH1) in the presence or absence of Exo. Lenti/Cont in the presence or absence of Exo was used as a control, respectively (B). The levels of nuclear p300-bound HIF-1α were determined with co-immunoprecipitation assay. The nuclear p300 was used as a loading control. The cellular levels of pan HIF-1α was examined. GAPDH was used as a loading control. **C and D**, A stable HeLa cell line expressing a HIF-1α-activated luciferase reporter gene was treated the same with panel A and B (n = 4). The relative activities of HIF-1α in the PBS, Exo, Exo/antimiR-Cont or Exo/antimiR-31 treated cells (C), and in the cells transduced with Lenti/Cont, Lenti/FIH1, Exo+Lenti/Cont or Exo+Lenti/FIH1 (D) were examined with luciferase assay. \**p*<0.05 and \*\**p*<0.01.

## Discussion

In spite of the significant advances in medical, interventional, and surgical therapies for ischemic diseases, the continued worldwide epidemic of these diseases urgently requires the development of innovative treatments. At this stage, an emerging microvesicle/exosome approach of therapeutic angiogenesis may present a promising outlook[31]. Here, we demonstrate that exosomal miR-31 enhances angiogenesis in ischemic animal models through an FIH1/HIF-1α signaling pathway.

PAD and ischemic heart disease represent two large groups of ischemic diseases and contribute to global disability and mortality. The premier characteristic of these ischemic diseases is the hypoxic microenvironment (ischemic hypoxia) induced by the blockage of blood vessels. Emerging evidence in exosome-based therapy indicates that exosomes promote therapeutic angiogenesis in ischemic tissues, although the underlying molecular mechanisms remain to be fully elucidated[32, 33]. In this study, we have demonstrated that miRNA in ASC exosomes contributes to exosome-induced angiogenesis in ischemic hindlimb and heart. Similar observations are reported in ischemic hindlimb by using exosomes from CD34^+^ cells isolated from peripheral blood[32], and in ischemic heart by using ASC-exosomes[34]. The former literature claims that a miR-126-3p is involved, whereas an overexpressing of miR-126 in exosomal donor cells is manipulated in the latter one. In our study, we not only take the advantage of ASCs as easy acquisition and less immunogenicity, but also pretreat ASCs with endothelial differentiation medium, without manipulation, to obtain much more exosomes enriched in proangiogenic miR-31.

Deficiency of miR-31 in plasma and endothelial progenitor cells from patients with coronary artery disease is observed. Intramuscular transplantation of the endothelial progenitor cells, upon restoration of miR-31, promotes angiogenesis in a rat HLI model[35]. These observations of proangiogenic miR-31 are in accordance with our results. Interestingly, in a rat MI model, up-regulation of cardiac miR-31 is observed. Subcutaneous administration of LNA (locked nucleic acid) inhibitor of miR-31 preserves cardiac function and decreases infarct size[36]. The different effects of miR-31 may be due to the administration of only an inhibitor, not cells or exosomes. FIH1 has been validated to be a direct target of miR-31 in vascular endothelial cells in our previous study[7] and in other cell types[18, 19]. Exosomal miR-135b from multiple myeloma cells enhances tube formation of endothelial cells and angiogenesis in transplanted plugs by targeting FIH1[37], which is consistent with our results that exosomal miR-31 from ASCs promotes angiogenesis in ischemic hindlimb and heart by targeting FIH1. Interestingly, administration of short hairpin RNA to inhibit FIH1 fails to show significant beneficial effect on a mouse MI model[14], which may be caused by a shorter half-life of short hairpin RNA in comparison with exosome-encapsulated miRNA.

Besides prolyl hydroxylases, which down-regulates HIF-1α stability via post-transnational modification under normoxic conditions, FIH1, as an asparaginyl hydroxylase, down-regulates HIF-1α transcriptional activity. In contrast, under hypoxic conditions hydroxylation of HIF-1α asparagine (N803) by FIH1 is diminished. This enables HIF-1α to bind with its co-activators, resulting in transcriptional activation of the target genes[13, 38]. In this study, we demonstrate that miR-31-enriched exosomes augment the binding of HIF-1α to p300 and enhance the transcriptional activity of HIF-1α, which are abrogated by overexpression of FIH1. To initiate the expression of proangiogenic factors is one of the most important roles of HIF-1α in ischemic diseases[11, 39]. Considering the complex composition in exosomes, the other miRNAs or proteins may also contribute the beneficial effects of exosomes on proangiogenesis. We recently reveal, for example, that miR-21 delivered by exosomes from hypoxic ASCs indirectly promote angiogenesis by polarizing macrophage to M2 phenotype[24]. According to the clinicaltrials.gov website, only one clinical trial that interventionally administers exosomes is completed so far, yet the results are still unpublished (NCT04276987, Phase I).

In summary, our results demonstrate that ASC exosomes promote angiogenesis in the mouse ischemic hindlimb and the ischemic heart via delivery of miR-31, which targets FIH1 in vascular endothelial cells and thus initiates HIF-1α transcriptional activity. These findings provide new insight into the mechanism underlying the application of exosomes from bench to bedside. As more mechanistic findings are uncovered, the emerging role of exosomes may promise a prospect of a novel exosome-based therapy for various diseases.

## Abbreviations

α-SMA: α-smooth muscle actin
antimiR-31: Lenti/anti-miR-31
antimiR-Cont: Lenti/anti-miR-Cont
ASC: adipose-derived stem cell
Exo: exosome
FIH1: factor inhibiting hypoxia-inducible factor-1
HIF-1α: hypoxia-inducible factor-1α
HMVEC: human microvascular endothelial cell
HLI: hindlimb ischemia
LAD: left anterior descending
LVESD: left ventricle (LV) end-systolic diameter
LVEDD: left ventricle end-diastolic diameter
LVFS%: percentage of left ventricle fractional shortening
MI: Myocardial infarction
PAD: peripheral artery disease

## Acknowledgments

We thank Dr. Ming-Bo Huang for the assistance with nanoparticle tracking analysis, Dr. Yan Xiao for the assistance with immunohistochemistry.

## REFERENCES

[1] N.D. Wong, Epidemiological studies of CHD and the evolution of preventive cardiology, Nat Rev Cardiol 11(5) (2014) 276–89.

[2] L. Zhao, T. Johnson, D. Liu, Therapeutic angiogenesis of adipose-derived stem cells for ischemic diseases, Stem Cell Res Ther 8(1) (2017) 125.

[3] J. Tongers, D.W. Losordo, U. Landmesser, Stem and progenitor cell-based therapy in ischaemic heart disease: promise, uncertainties, and challenges, Eur Heart J 32(10) (2011) 1197–206.

[4] T. Johnson, L. Zhao, G. Manuel, H. Taylor, D. Liu, Approaches to therapeutic angiogenesis for ischemic heart disease, J Mol Med (Berl) 97(2) (2019) 141–151.

[5] R. Kalluri, V.S. LeBleu, The biology, function, and biomedical applications of exosomes, Science 367(6478) (2020).

[6] R. Madonna, Y.J. Geng, R. De Caterina, Adipose tissue-derived stem cells: characterization and potential for cardiovascular repair, Arterioscler Thromb Vasc Biol 29(11) (2009) 1723–9.

[7] T. Kang, T.M. Jones, C. Naddell, M. Bacanamwo, J.W. Calvert, W.E. Thompson, V.C. Bond, Y.E. Chen, D. Liu, Adipose-Derived Stem Cells Induce Angiogenesis via Microvesicle Transport of miRNA-31, Stem Cells Transl Med 5(4) (2016) 440–50.

[8] X. Liang, L. Zhang, S. Wang, Q. Han, R.C. Zhao, Exosomes secreted by mesenchymal stem cells promote endothelial cell angiogenesis by transferring miR-125a, J Cell Sci 129(11) (2016) 2182–9.

[9] F. Figliolini, A. Ranghino, C. Grange, M. Cedrino, M. Tapparo, C. Cavallari, A. Rossi, G. Togliatto, S. Femmino, M.V. Gugliuzza, G. Camussi, M.F. Brizzi, Extracellular Vesicles From Adipose Stem Cells Prevent Muscle Damage and Inflammation in a Mouse Model of Hind Limb Ischemia: Role of Neuregulin-1, Arterioscler Thromb Vasc Biol 40(1) (2020) 239–254.

[10] Q. Lv, J. Deng, Y. Chen, Y. Wang, B. Liu, J. Liu, Engineered Human Adipose Stem-Cell-Derived Exosomes Loaded with miR-21-5p to Promote Diabetic Cutaneous Wound Healing, Mol Pharm 17(5) (2020) 1723–1733.

[11] S. Rey, G.L. Semenza, Hypoxia-inducible factor-1-dependent mechanisms of vascularization and vascular remodelling, Cardiovasc Res 86(2) (2010) 236–42.

[12] A. Zimna, M. Kurpisz, Hypoxia-Inducible Factor-1 in Physiological and Pathophysiological Angiogenesis: Applications and Therapies, Biomed Res Int 2015 (2015) 549412.

[13] P.C. Mahon, K. Hirota, G.L. Semenza, FIH-1: a novel protein that interacts with HIF-1alpha and VHL to mediate repression of HIF-1 transcriptional activity, Genes Dev 15(20) (2001) 2675–86.

[14] M. Huang, P. Nguyen, F. Jia, S. Hu, Y. Gong, P.E. de Almeida, L. Wang, D. Nag, M.A. Kay, A.J. Giaccia, R.C. Robbins, J.C. Wu, Double knockdown of prolyl hydroxylase and factor-inhibiting hypoxia-inducible factor with nonviral minicircle gene therapy enhances stem cell mobilization and angiogenesis after myocardial infarction, Circulation 124(11 Suppl) (2011) S46–54.

[15] J.H. So, J.D. Kim, K.W. Yoo, H.T. Kim, S.H. Jung, J.H. Choi, M.S. Lee, S.W. Jin, C.H. Kim, FIH-1, a novel interactor of mindbomb, functions as an essential anti-angiogenic factor during zebrafish vascular development, PLoS One 9(10) (2014) e109517.

[16] R.A. Boon, S. Dimmeler, MicroRNAs in myocardial infarction, Nat Rev Cardiol 12(3) (2015) 135–42.

[17] D. Kir, E. Schnettler, S. Modi, S. Ramakrishnan, Regulation of angiogenesis by microRNAs in cardiovascular diseases, Angiogenesis 21(4) (2018) 699–710.

[18] H. Peng, N. Kaplan, R.B. Hamanaka, J. Katsnelson, H. Blatt, W. Yang, L. Hao, P.J. Bryar, R.S. Johnson, S. Getsios, N.S. Chandel, R.M. Lavker, microRNA-31/factor-inhibiting hypoxia-inducible factor 1 nexus regulates keratinocyte differentiation, Proc Natl Acad Sci U S A 109(35) (2012) 14030–4.

[19] B. Zhu, X. Cao, W. Zhang, G. Pan, Q. Yi, W. Zhong, D. Yan, MicroRNA-31-5p enhances the Warburg effect via targeting FIH, Faseb j 33(1) (2019) 545–556.

[20] W.D. Gray, K.M. French, S. Ghosh-Choudhary, J.T. Maxwell, M.E. Brown, M.O. Platt, C.D. Searles, M.E. Davis, Identification of therapeutic covariant microRNA clusters in hypoxia-treated cardiac progenitor cell exosomes using systems biology, Circ Res 116(2) (2015) 255–63.

[21] U. Agarwal, A. George, S. Bhutani, S. Ghosh-Choudhary, J.T. Maxwell, M.E. Brown, Y. Mehta, M.O. Platt, Y. Liang, S. Sahoo, M.E. Davis, Experimental, Systems, and Computational Approaches to Understanding the MicroRNA-Mediated Reparative Potential of Cardiac Progenitor Cell-Derived Exosomes From Pediatric Patients, Circ Res 120(4) (2017) 701–712.

[22] M. Chandy, J.W. Rhee, M.O. Ozen, D.R. Williams, L. Pepic, C. Liu, H. Zhang, J. Malisa, E. Lau, U. Demirci, J.C. Wu, Atlas of Exosomal microRNAs Secreted From Human iPSC-Derived Cardiac Cell Types, Circulation 142(18) (2020) 1794–1796.

[23] D. Liu, Y. Lin, T. Kang, B. Huang, W. Xu, M. Garcia-Barrio, M. Olatinwo, R. Matthews, Y.E. Chen, W.E. Thompson, Mitochondrial dysfunction and adipogenic reduction by prohibitin silencing in 3T3-L1 cells, PLoS One 7(3) (2012) e34315.

[24] D. Zhu, T.K. Johnson, Y. Wang, M. Thomas, K. Huynh, Q. Yang, V.C. Bond, Y.E. Chen, D. Liu, Macrophage M2 polarization induced by exosomes from adipose-derived stem cells contributes to the exosomal proangiogenic effect on mouse ischemic hindlimb, Stem Cell Res Ther 11(1) (2020) 162.

[25] E. Gao, Y.H. Lei, X. Shang, Z.M. Huang, L. Zuo, M. Boucher, Q. Fan, J.K. Chuprun, X.L. Ma, W.J. Koch, A novel and efficient model of coronary artery ligation and myocardial infarction in the mouse, Circ Res 107(12) (2010) 1445–53.

[26] J. Takagawa, Y. Zhang, M.L. Wong, R.E. Sievers, N.K. Kapasi, Y. Wang, Y. Yeghiazarians, R.J. Lee, W. Grossman, M.L. Springer, Myocardial infarct size measurement in the mouse chronic infarction model: comparison of area- and length-based approaches, J Appl Physiol (1985) 102(6) (2007) 2104–11.

[27] M. Colombo, G. Raposo, C. Thery, Biogenesis, secretion, and intercellular interactions of exosomes and other extracellular vesicles, Annu Rev Cell Dev Biol 30 (2014) 255–89.

[28] C. Thery, et al., Minimal information for studies of extracellular vesicles 2018 (MISEV2018): a position statement of the International Society for Extracellular Vesicles and update of the MISEV2014 guidelines, J Extracell Vesicles 7(1) (2018) 1535750.

[29] J.P. Cooke, S. Meng, Vascular Regeneration in Peripheral Artery Disease, Arterioscler Thromb Vasc Biol 40(7) (2020) 1627–1634.

[30] D. Lando, D.J. Peet, J.J. Gorman, D.A. Whelan, M.L. Whitelaw, R.K. Bruick, FIH-1 is an asparaginyl hydroxylase enzyme that regulates the transcriptional activity of hypoxia-inducible factor, Genes Dev 16(12) (2002) 1466–71.

[31] D. Todorova, S. Simoncini, R. Lacroix, F. Sabatier, F. Dignat-George, Extracellular Vesicles in Angiogenesis, Circ Res 120(10) (2017) 1658–1673.

[32] P. Mathiyalagan, Y. Liang, D. Kim, S. Misener, T. Thorne, C.E. Kamide, E. Klyachko, D.W. Losordo, R.J. Hajjar, S. Sahoo, Angiogenic Mechanisms of Human CD34(+) Stem Cell Exosomes in the Repair of Ischemic Hindlimb, Circ Res 120(9) (2017) 1466–1476.

[33] A. Robson, Exosomes improve myocardial recovery after infarction, Nat Rev Cardiol 17(12) (2020) 758.

[34] Q. Luo, D. Guo, G. Liu, G. Chen, M. Hang, M. Jin, Exosomes from MiR-126-Overexpressing Adscs Are Therapeutic in Relieving Acute Myocardial Ischaemic Injury, Cell Physiol Biochem 44(6) (2017) 2105–2116.

[35] H.W. Wang, T.S. Huang, H.H. Lo, P.H. Huang, C.C. Lin, S.J. Chang, K.H. Liao, C.H. Tsai, C.H. Chan, C.F. Tsai, Y.C. Cheng, Y.L. Chiu, T.N. Tsai, C.C. Cheng, S.M. Cheng, Deficiency of the microRNA-31-microRNA-720 pathway in the plasma and endothelial progenitor cells from patients with coronary artery disease, Arterioscler Thromb Vasc Biol 34(4) (2014) 857–69.

[36] E.C. Martinez, S. Lilyanna, P. Wang, L.A. Vardy, X. Jiang, A. Armugam, K. Jeyaseelan, A.M. Richards, MicroRNA-31 promotes adverse cardiac remodeling and dysfunction in ischemic heart disease, J Mol Cell Cardiol 112 (2017) 27–39.

[37] T. Umezu, H. Tadokoro, K. Azuma, S. Yoshizawa, K. Ohyashiki, J.H. Ohyashiki, Exosomal miR-135b shed from hypoxic multiple myeloma cells enhances angiogenesis by targeting factor-inhibiting HIF-1, Blood 124(25) (2014) 3748–57.

[38] A. Weidemann, R.S. Johnson, Biology of HIF-1alpha, Cell Death Differ 15(4) (2008) 621–7.

[39] T. Bishop, P.J. Ratcliffe, HIF hydroxylase pathways in cardiovascular physiology and medicine, Circ Res 117(1) (2015) 65–79.

