## Supplemental data for "Exosomes from adipose-derived stem cells alleviate myocardial infarction via microRNA-31/FIH1/HIF-1α pathway"

### Slide 1
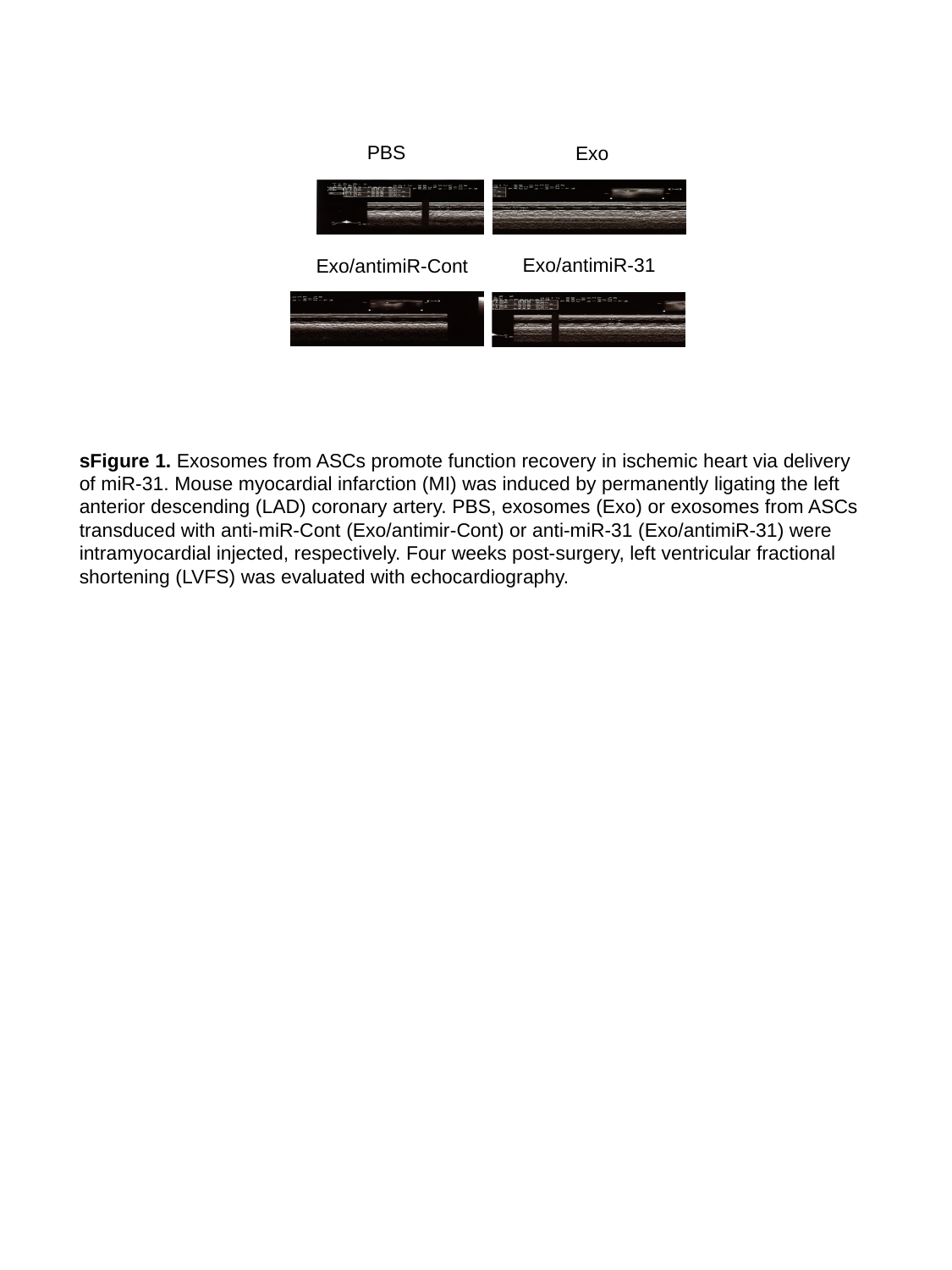

PBS
Exo
Exo/antimiR-31
Exo/antimiR-Cont
sFigure 1. Exosomes from ASCs promote function recovery in ischemic heart via delivery of miR-31. Mouse myocardial infarction (MI) was induced by permanently ligating the left anterior descending (LAD) coronary artery. PBS, exosomes (Exo) or exosomes from ASCs transduced with anti-miR-Cont (Exo/antimir-Cont) or anti-miR-31 (Exo/antimiR-31) were intramyocardial injected, respectively. Four weeks post-surgery, left ventricular fractional shortening (LVFS) was evaluated with echocardiography.

### Slide 2
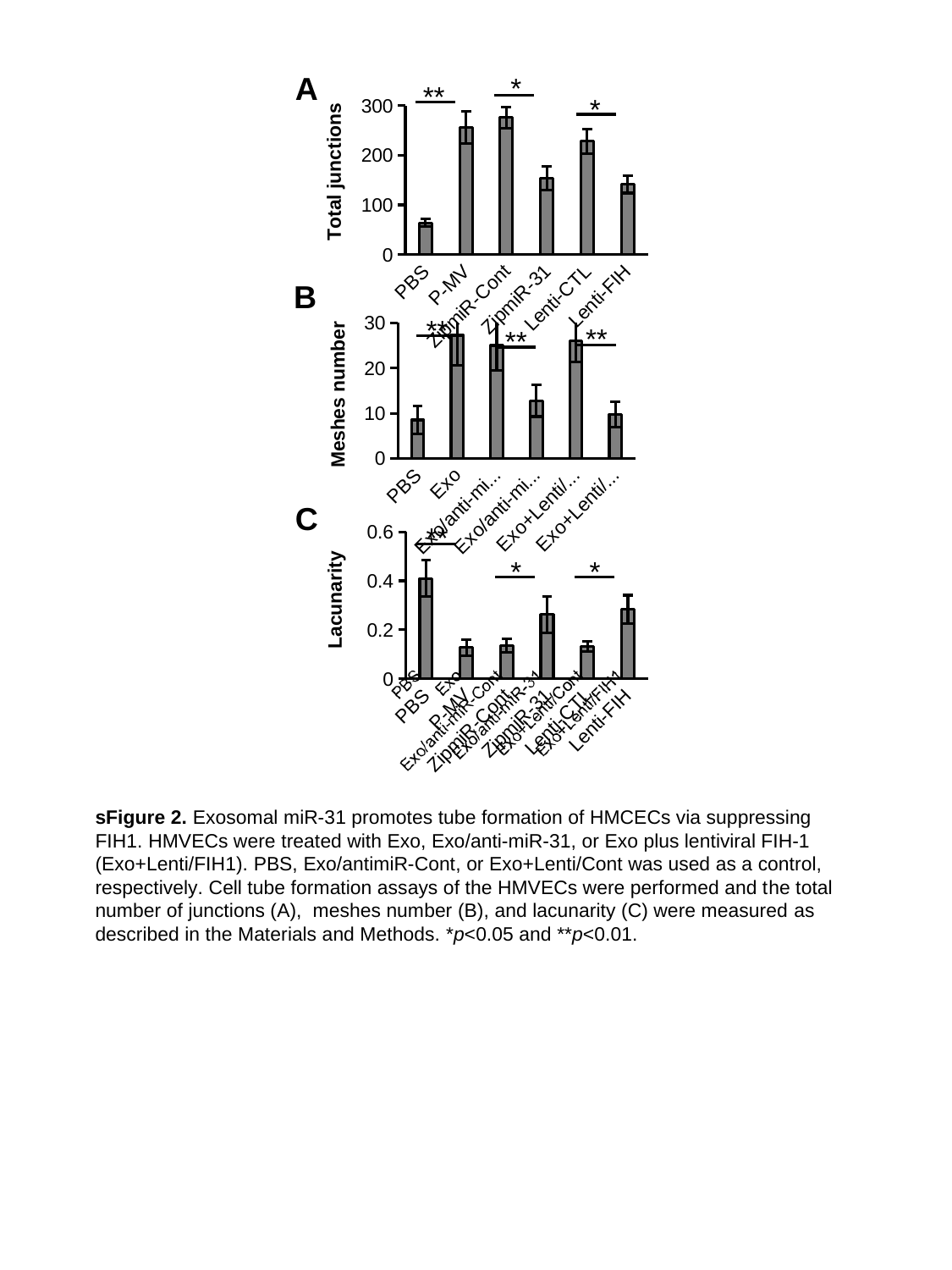

A
*
**
*
#### Chart
| Category | |
|---|---|
| PBS | 63.75 |
| P-MV | 255.75 |
| ZipmiR-Cont | 276.0 |
| ZipmiR-31 | 153.5 |
| Lenti-CTL | 228.0 |
| Lenti-FIH | 141.5 |B
#### Chart
| Category | |
|---|---|
| PBS | 8.5 |
| Exo | 27.25 |
| Exo/anti-miR-Cont | 25.0 |
| Exo/anti-miR-31 | 12.75 |
| Exo+Lenti/Cont | 26.0 |
| Exo+Lenti/FIH1 | 9.75 |**
**
**
C
**
#### Chart
| Category | |
|---|---|
| PBS | 0.41 |
| P-MV | 0.1275 |
| ZipmiR-Cont | 0.135 |
| ZipmiR-31 | 0.2625 |
| Lenti-CTL | 0.1325 |
| Lenti-FIH | 0.2825 |*
*
sFigure 2. Exosomal miR-31 promotes tube formation of HMCECs via suppressing FIH1. HMVECs were treated with Exo, Exo/anti-miR-31, or Exo plus lentiviral FIH-1 (Exo+Lenti/FIH1). PBS, Exo/antimiR-Cont, or Exo+Lenti/Cont was used as a control, respectively. Cell tube formation assays of the HMVECs were performed and the total number of junctions (A),  meshes number (B), and lacunarity (C) were measured as described in the Materials and Methods. *p<0.05 and **p<0.01.

### Slide 3
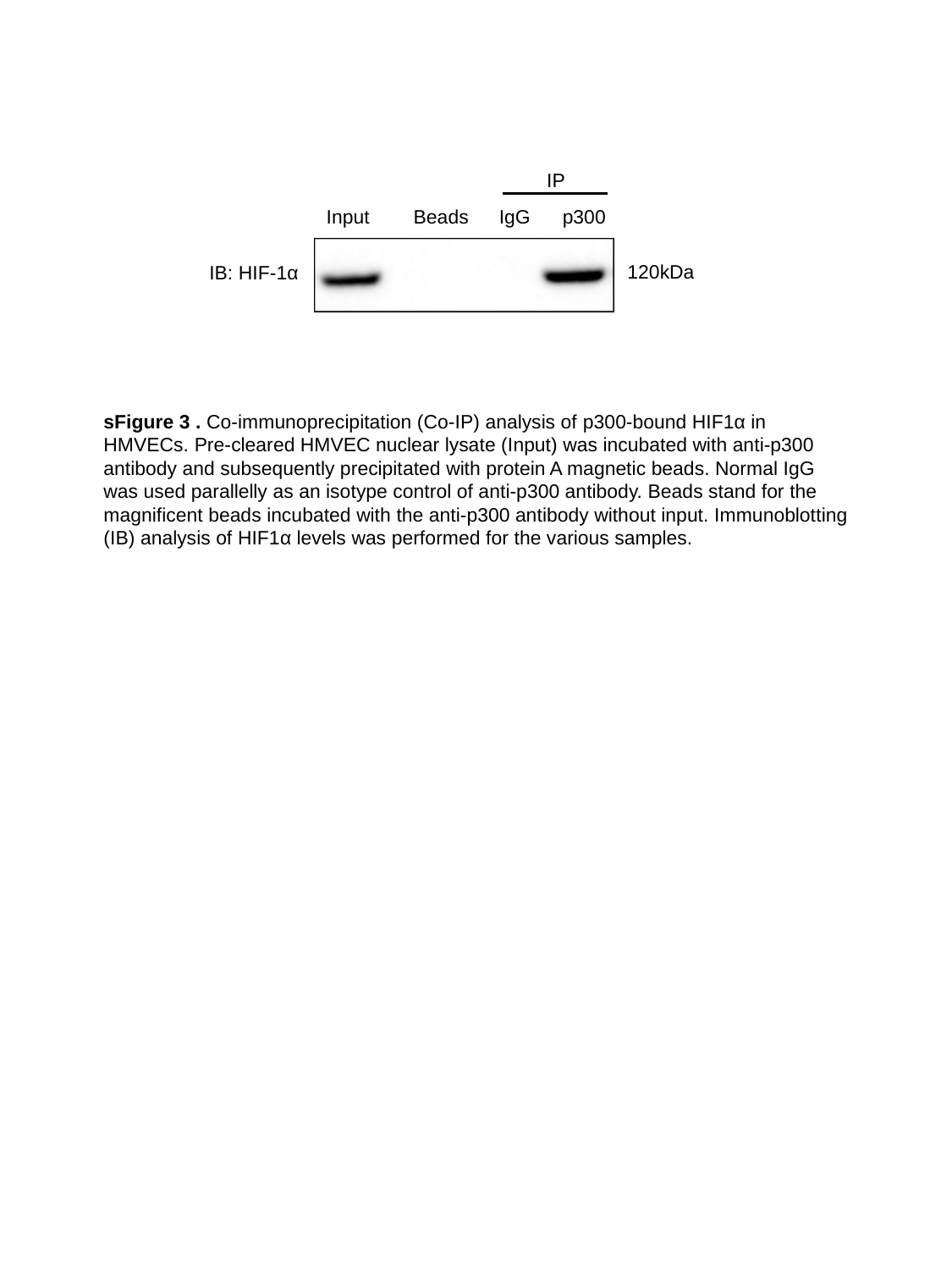

IP
Input
IgG
p300
Beads
120kDa
IB: HIF-1α
sFigure 3 . Co-immunoprecipitation (Co-IP) analysis of p300-bound HIF1α in HMVECs. Pre-cleared HMVEC nuclear lysate (Input) was incubated with anti-p300 antibody and subsequently precipitated with protein A magnetic beads. Normal IgG was used parallelly as an isotype control of anti-p300 antibody. Beads stand for the magnificent beads incubated with the anti-p300 antibody without input. Immunoblotting (IB) analysis of HIF1α levels was performed for the various samples.

### Slide 4
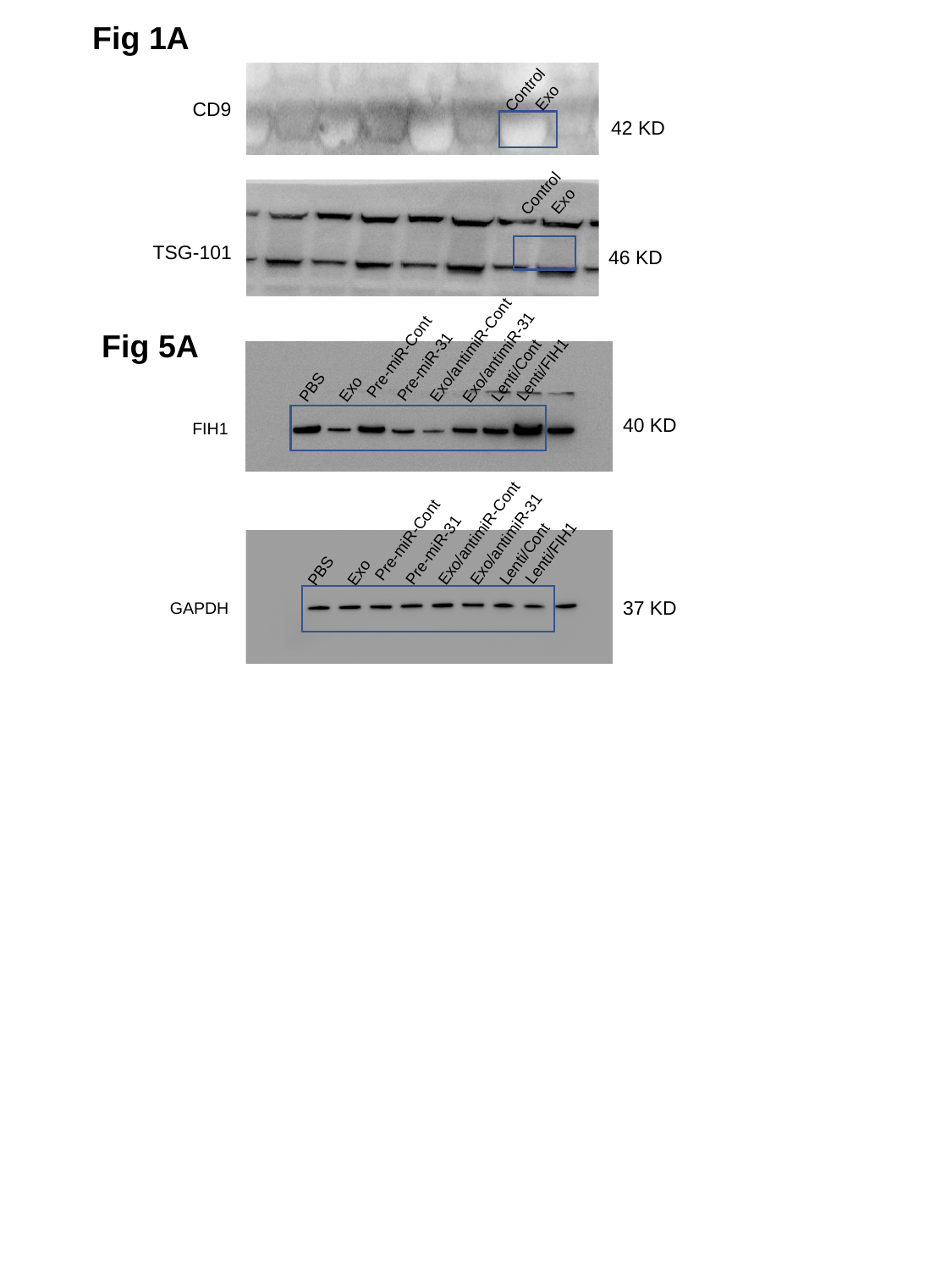

Fig 1A
Control
Exo
CD9
42 KD
Control
Exo
TSG-101
46 KD
Fig 5A
Pre-miR-31
Exo/antimiR-Cont
Exo/antimiR-31
Pre-miR-Cont
Lenti/Cont
Lenti/FIH1
Exo
PBS
40 KD
FIH1
Pre-miR-31
Exo/antimiR-Cont
Exo/antimiR-31
Pre-miR-Cont
Lenti/Cont
Lenti/FIH1
Exo
PBS
37 KD
GAPDH

### Slide 5
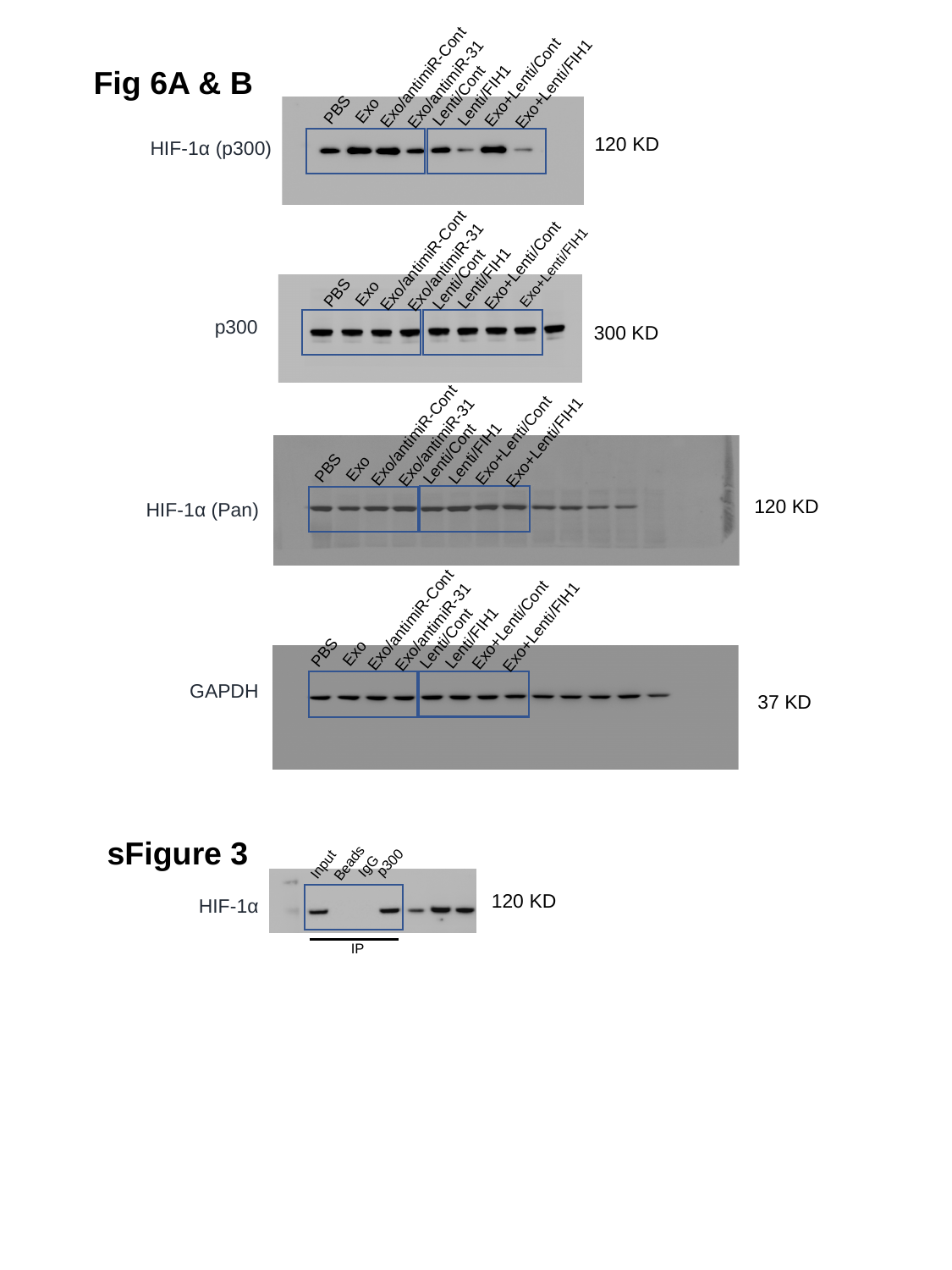

Fig 6A & B
Exo/antimiR-Cont
Exo+Lenti/Cont
Exo/antimiR-31
Exo+Lenti/FIH1
Lenti/FIH1
Lenti/Cont
PBS
Exo
120 KD
HIF-1α (p300)
Exo/antimiR-Cont
Exo+Lenti/Cont
Exo/antimiR-31
Exo+Lenti/FIH1
Lenti/FIH1
Lenti/Cont
PBS
Exo
p300
300 KD
Exo/antimiR-Cont
Exo+Lenti/Cont
Exo/antimiR-31
Exo+Lenti/FIH1
Lenti/FIH1
Lenti/Cont
PBS
Exo
120 KD
HIF-1α (Pan)
Exo/antimiR-Cont
Exo+Lenti/Cont
Exo/antimiR-31
Exo+Lenti/FIH1
Lenti/FIH1
Lenti/Cont
PBS
Exo
GAPDH
37 KD
sFigure 3
p300
Beads
Input
IgG
120 KD
HIF-1α
IP
